# Droplet on Demand Tape Drive and XES Prototypes for Time-Resolved Serial Crystallography at VMXi, Diamond Light Source

**DOI:** 10.1101/2025.11.19.689295

**Authors:** Pierre Aller, Juan Sanchez-Weatherby, Abbey Telfer, Robert Bosman, Nicholas Devenish, Philip Hinchliffe, Sam Horrell, Jophy Ip, Richard Littlewood, Andrew Male, Eva Giminez Navarro, Urszula Neuman, Jos J. A. G. Kamps, David Omar, Laura Parkinson, Muthraj Pandi, Nico Rubies, James Sandy, Anastasiia Shilova, James Spencer, Jonathan Spiers, John P. Sutter, Amy J. Thompson, Catherine L. Tooke, Ben Williams, Tiankun Zhou, Michael A. Hough, Allen M. Orville

**Affiliations:** Diamond Light Source Ltd., Harwell Science and Innovation Campus, Didcot, OX11 0DE, UK; Research Complex at Harwell, Rutherford Appleton Laboratory, Didcot, OX11 0FA, UK; School of Biochemistry and Cellular and Molecular Medicine, Biomedical Sciences Building, University of Bristol, Bristol BS8□1TD, UK; Department of Biology and Biochemistry, 4 South, University of Bath, Bath BA2 7AY, UK; University Medical Center Hamburg-Eppendorf (UKE), Hamburg, Germany; Department of Life Sciences, Sir Ernst Chain Building, Imperial College London, London, SW7 2AZ, UK; European Synchrotron Radiation Facility (ESRF), Grenoble, France; Bruker UK Limited; University of Oxford, Department of Statistics 24-29 St Giles, Oxford OX1 3LB, UK; Oxford Physics Microstructure Detector (OPMD) laboratory, Department of Physics, University of Oxford, Park Road, Oxford OX1 3PU, UK

**Keywords:** Serial synchrotron crystallography (SSX), droplet on demand tape drive, X-ray emission spectroscopy (XES), metalloenzyme

## Abstract

Time resolved X-ray crystallography is experiencing a resurgence, in part, because of serial methods that readily allow scientists to create stop-motion movies of macromolecular function of photoactivation, enzyme catalysed reactions, and ligand-induced conformational changes triggering further downstream signalling events. While some reactions can be initiated with light, either naturally or using photocaged compounds, a more generally applicable approach is to mix microcrystals with reagents at varying time points prior to exposure to the X-ray beam. A powerful approach has been to combine droplet on demand ‘tape drive’ sample delivery with X-ray emission spectroscopy (XES) that correlates atomic structure with electronic states of metal ions within the sample. To our knowledge, such a combined methodology has not been deployed previously at a synchrotron beamline and has been restricted to XFELs. Here we describe prototype experiments along the development pathway to a combined droplet on demand diffraction and XES system at the microfocus beamline VMXi at Diamond Light Source. We demonstrate the collection of a high-quality serial diffraction data set from microcrystals within hundreds of picolitre-volume droplets deposited on a moving tape. In separate experiments at VMXi, we collected XES data from microcrystals of a copper enzyme delivered using a high viscosity extruder. Together, these results demonstrate the feasibility of combined droplet on demand serial crystallography and XES experiments using a third-generation synchrotron beamline; project work currently underway at Diamond Light Source.

**Synopsis:** We describe proof of concept experiments towards correlated serial crystallography (SSX) and X-ray emission spectroscopy (XES) from microcrystals at a microfocus synchrotron beamline. A droplet on demand tape drive system delivers microcrystals to the beam within well-separated, hundreds of picolitre-volume droplets while XES allows validation of redox states of metals within protein crystals.

## 1. Introduction

The emergence of X-ray Free Electron Laser (XFEL) facilities motivated development of serial femtosecond crystallography (SFX) data collection strategies, including many sample delivery methods that are amenable to time-resolved studies (Barends et al., 2022; Branden & Neutze, 2021; Chapman, 2019). Serial crystallography is becoming an increasingly popular tool for structural biologists, especially at synchrotrons where beamtime is typically more readily available than at XFEL sources (Aller & Orville, 2021; Orville, 2020; Pearson & Mehrabi, 2020). Third and fourth generation synchrotron beamlines, with high flux and microbeam capabilities, are particularly well suited for serial crystallography (Jaho et al., 2024; Mehrabi, Schulz, Dsouza, et al., 2019). The advent of diffraction limited lattices (so called “fourth generation” synchrotrons), enables even smaller beam sizes and significantly greater flux densities. Several beamlines are now fully dedicated to performing serial crystallography, including ID29 at ESRF-EBS (Orlans et al., 2025), T-REXX at PETRA III (Mehrabi, Schulz, Agthe, et al., 2019) and MicroMAX at MAX IV (Gonzalez et al., 2025). At Diamond Light Source, two macromolecular crystallography beamlines are particularly suitable for serial crystallography: I24 and VMXi. The micro-focus beamline I24 has developed a world leading, flexible sample mounting system that enables rapid switching between a rotation and serial sample delivery setup. This permits the beamline to operate different data-collection modes, either a “pin” mode or “serial” mode. Furthermore, I24 can use different serial sample delivery setups either a fixed target or a high viscosity extruder (Birch et al., 2023; Jaho et al., 2024). On the other hand, VMXi was designed for *in-situ* room temperature data collection from crystals within crystallisation plates. Moreover, VMXi is also well suited for serial data collection since it incorporates a multi-layer monochromator (DMM) producing a broad energy bandpass (∼5 × 10□^3^ ΔE/E), resulting in high flux (>2 × 10^13^ ph/s at 16 keV) with a microfocus pink beam (10 ×10 µm) (Sanchez-Weatherby et al., 2019; Sandy et al., 2024). The characteristics of VMXi make it well suited for serial crystallography, in terms of flux, beam size and detector repetition rate (Eiger2X 4M, running in continuous mode at 500 Hz).

Time-resolved crystallography studies typically require interpreting electron density maps that represent mixed intermediate populations. This often presents different possible reaction intermediates at a certain time-point, which may or may not be easy to resolve from the electron density alone. Consequently, complementary spectroscopic methods are incredibly valuable for time-resolved studies since they can provide additional orthogonal data to help resolve ambiguities (Wilmot & Pearson, 2002). When studying metalloproteins it is possible to combine X-ray spectroscopy methods with X-ray crystallography. This can provide a critical advantage especially if/when the X-ray probed region simultaneously yields diffraction and spectroscopic information because it removes uncertainty arising from data collection conditions (*i*.*e*. an offset in time, or over differing exposures, *etc*.). An important caveat is that spectroscopy “sees” all the metal ions and may not differentiate between ordered and disordered species, whereas X-ray diffraction and resulting electron density maps reveal only ordered atoms (Einsle et al., 2007; Sauter et al., 2020; Zhang et al., 2013). Combining serial crystallography with X-ray emission spectroscopy (XES) is therefore an elegant strategy for obtaining well-correlated types of data from the same sample and X-ray pulse/exposure (Fuller et al., 2017; Kamps et al., 2024). Thus, XES data can help validate the interpretation of electron density maps, for catalytically active metal centres within metalloenzymes in time-resolved experiments.

An effective way to collect XES while simultaneously collecting diffraction images uses a von Hamos geometry spectrometer. Developed for XFEL applications (Alonso-Mori, Kern, Gildea, et al., 2012; Alonso-Mori, Kern, Sokaras, et al., 2012; Kern et al., 2013), a consortium including ourselves have demonstrated the drop on tape method (DoT) combining XES and diffraction experiments (Butryn et al., 2021; Kamps et al., 2024; Rabe et al., 2021). The DoT strategy, originally developed at a synchrotron facility as described previously (Roessler et al., 2013), is now predominantly employed at XFELs (Fuller et al., 2017). The main advantages of the DoT sample delivery system are i) the low sample consumption using nanolitre sample droplets ejected on demand and synchronized with the X-ray pulse, unlike jet techniques which also dispense sample between X-ray pulses, ii) versatility of the solid support allowing diverse reaction initiation methods such as turbulent mixing, gas diffusion, and light illumination (Kamps et al., 2024) and iii) the possibility to exploit anaerobic conditions (Lebrette et al., 2023; Rabe et al., 2021). The DoT system developed by the consortium at LCLS uses 2-5 nanolitre size droplets and evolved into a complicated to install and operate data collection strategy requiring significant support from the consortium. We have long noted that working with smaller droplets would significantly reduce sample consumption and improve the temporal resolution of time-resolved data.

Meanwhile, alternative tape-based sample delivery methods—such as those using capillaries to produce continuous sample streams—are being actively developed and used at synchrotron sources (Kamps et al., 2024). The Center for Free-Electron Laser Science (CFEL), Hamburg, developed a simpler tape drive (CFEL TapeDrive 2.0) operating as a reel to reel and using microfluidics to mix microcrystal slurry with ligand or substrate before depositing a streak of the mixture onto the moving Kapton^®^ tape (Beyerlein et al., 2017; Henkel et al., 2023; Zielinski et al., 2022). This tape drive is used at P11 (Petra III) and MicroMAX (Max IV); a similar tape drive exists at ID29 (ESRF-EBS). The main advantage of these systems is the ease of use, whereas a limitation is the lack of versatility for implementation of complementary methods.

K-shell XES is an element selective technique which probes an atom’s electronic properties (*i*.*e*. redox and spin states) by measuring the energy dependence of emitted photons when an outer shell electron fills the K-shell hole created by the incident X-ray beam. (Cutsail & DeBeer, 2022; Glatzel & Bergmann, 2005). For 3d transition metals the three most relevant emission lines are the Kα_1,2_ (2p-1s), Kβ_1,3_ (3p-1s) and valence to core (Kβ_2,5_) in decreasing order of their signal intensity. The Kα_1,2_ doublet is a reporter of oxidation state while Kβ_1,3_ is sensitive to metal-ligand covalency, spin state and oxidation state. The weakest (valence to core) signal can offer insights into the metal ion coordination environment providing information on binding and the nature of valence molecular orbitals, but is also highly challenging to measure accurately due to its weak signal.

Combining time-resolved XES with serial crystallography has been successful in monitoring metal centres where the oxidation state correlates with enzyme reaction intermediates. In the iron and oxygen dependent isopenicillin N-synthase (IPNS), time resolved XES with high resolution SFX was measured from microcrystals co-crystallised with the tripeptide substrate under anaerobic conditions and the reaction initiated with exposure to 100% O_2_ gas. This identified structural intermediates and conformational changes triggered by oxygen binding. The correlated XES monitored the width of the Fe Kα_1,2_ peak and was consistent with the formation of Fe(III) valence state in the putative iron-bound superoxide intermediate and the associated structural changes throughout the enzyme that were revealed in the SFX data (Rabe et al., 2021). In the case of methane monooxygenase, Fe Kα_1,2_ XES data were used to verify Fe(III) -vs-Fe(II) status of the dinuclear metal centre that is very susceptible to photoreduction by X-ray exposure (Srinivas et al., 2020). Time-resolved SFX and XES data from the P450 enzyme CYP121 from *Mycobacterium tuberculosis* catalysing a shunt reaction with peracetic acid revealed a ferric-hydroperoxo intermediate in the 200 ms timepoint data that is fully correlated with its 1.85 Å resolution atomic structure (Nguyen et al., 2023). Finally, Mn Kβ_1,3_ tr-XES and tr-SFX was used to follow the S1 → S2 → S3 → [S4] → S0 transitions of the active oxygen evolving complex in photosystem II triggered by pump-probe reaction initiation strategies (Ibrahim et al., 2020).

Given these advantages, an important aspect for our tape drive development programme is the integration with XES. A challenge when working on metal containing proteins is the possibility of site-specific X-ray radiation induced perturbations, which can beneficially promote mechanistically relevant reactions, *e*.*g*. in redox proteins and enzymes, or lead to artificial redox state changes and experimental artefacts that may compromise the experimental outcome (Beitlich et al., 2007; Chreifi et al., 2016; Hajdu et al., 2000). Serial crystallography helps to mitigate the X-ray induced artifacts in the sample by spreading the X-ray dose over typically thousands of microcrystals. Additionally, it can be challenging to identify conditions and timepoints where catalytic intermediates have built up to high occupancy. Spectroscopic measurements from crystals provide complementary information, particularly when experiments are fully correlated such that different data types emerge from the same microcrystal volume. The additional data aids the interpretation of the electron density maps and refinement(s) of atomic model(s). Our goal is to exploit XES signal at both synchrotron and XFEL beamlines using the same instrumentation at both types of facilities.

Here we describe results from proof-of-concept experiments with prototype components for the DoT with XES instrument under development at VMXi at Diamond Light Source. Serial crystallographic data measured from protein microcrystals within hundreds pL droplets (≤ 500 pL) on a moving Kapton^®^ tape are described. A second experiment reports XES results for copper Kα_1,2_ measurements obtained from protein microcrystals using a four-analyser-crystal von Hamos spectrometer coupled with sample delivery using a viscous extruder. These results indicate the feasibility of carrying out this combined approach at a microfocus synchrotron beamline.

Our key design requirements were to keep the tape drive simple to use, transportable, and compact due to the space constraints at VMXi with the goal of portability to enable experiments at other beamlines. The prototype tape drive at VMXi demonstrated the ability to successfully obtain good quality serial crystallography data at high resolution, from microcrystals of the serine β-lactamase CTX-M-15. The von Hamos XES spectrometer prototype allowed representative measurements for Fe and Cu Kα_1,2_ spectra of salts and then copper nitrite reductase (CuNiR) microcrystals. These results demonstrate that combined DoT/XES data collection is feasible at a third-generation synchrotron source, with further improvements expected from the diffraction limited machine lattice at Diamond-II (https://www.diamond.ac.uk/Diamond-II.html), or at XFELs.

## 2. Methods

### 2.1.1. Sample preparation for tape drive experiments

Recombinant His-tagged CTX-M-15 was expressed and purified as previously described (Tooke et al., 2019). CTX-M-15 microcrystal slurries were generated as previously described (Butryn et al., 2021), using crystal seed stock that was generated by crushing macro-crystals grown by sitting drop vapour diffusion (Tooke et al., 2019); protein (5 µL) was mixed with crystallisation solution [5 µL; 2.0 M (NH_4_)_2_SO_4_, 0.1 M Tris pH 8] and seed (5 µL), and equilibrated against crystallisation solution (500 µL) in 24-well sitting drop plates (Hampton). Rod-shaped crystals grew within 24 h, with a maximum width of 3-8 µm and length around 8 µm. Crystal density was ∼1 x 10^8^ crystals mL^−1^ as measured using a TC20 automated cell counter (BioRad) or a counting chamber (Neubauer).

### 2.1.2. VMXi instrumentation setup and installation of tape drive prototype

The tape drive prototype consisted of a motor and tape, wash-dry mechanism, and sample ejection system (Fig. 1 and Fig. S1). First, the system was installed and aligned to the VMXi X-ray beam path. The typical layout of the beamline has been described previously (Sanchez-Weatherby et al., 2019; Sandy et al., 2024). An aluminium breadboard (MB2060/M, Thorlabs) mounted on MiniTec framework above the VMXi goniometer enabled the installation of the prototype tape drive as shown in Fig. 1a. In standard VMXi operation, the X-ray beam is focused on the rotation axis of the beamline goniometer (Fig. S2a). Due to the space requirements of the tape drive system, the sample interaction point for the experiments described here was instead placed 250 mm downstream from the standard position by modifying the settings of the microfocus bimorph mirrors (Fig. S2b). An extension tube was designed to visualise the beam at the newly created reference point using a drilled lens (OPTEM 35-00-02-000, 4x magnification and 50 mm working distance) to allow the X-ray beam to pass through. In this way the X-ray beam could be focused onto 50 µm YAG:Ce (Yttrium aluminium garnet (YAG) activated by cerium; Crytur, Czech Republic) X-ray imaging screen at the new downstream sample position and the beamline on-axis viewing system could resolve the new focal point. The X-ray focal point and the interaction region of the tape drive system was aligned by referencing with the on-axis viewing camera. A second camera was placed orthogonal to the sample interaction region that allowed continuous monitoring of the vertical and horizontal droplet position while collecting diffraction data.

**Figure 1.**
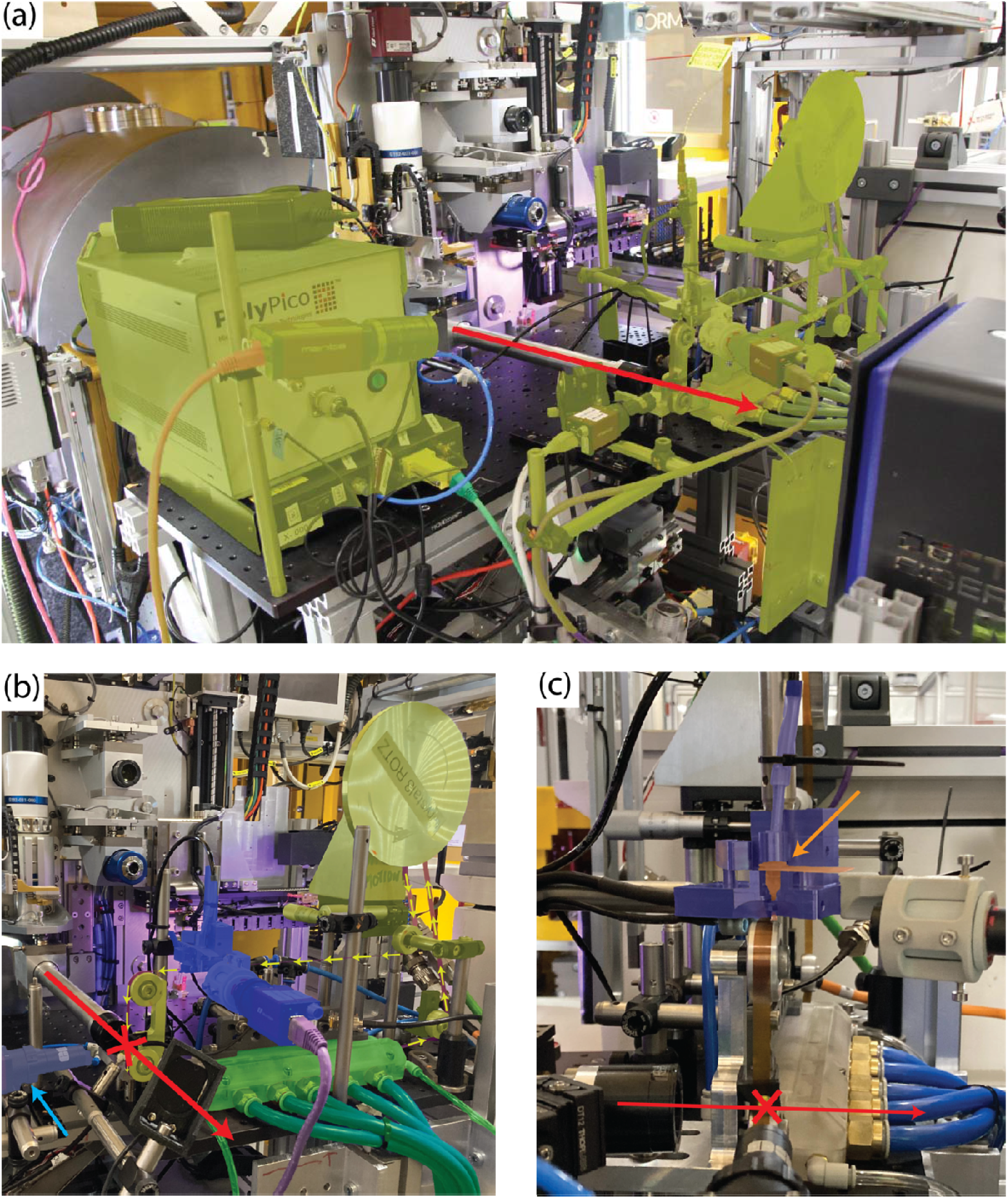
Tape drive prototype at VMXi. (a) Overall view of the installation of the tape drive prototype at the VMXi beamline. All the tape drive components are highlighted in yellow. The beam direction is given by the red arrow. (b) Focus on the tape drive prototype: mechanical components of the tape drive (motor and pulleys) are highlighted in yellow, the Kapton^®^ tape is highlighted in pink (with yellow arrows indicating the direction), the clean/dry unit highlighted in green, the piezoelectric injector (PEI) unit (camera, PolyPico mini-head and LED) highlighted in blue and orthogonal view camera highlighted in light blue (blue arrow). (c) Close-up view of the PolyPico mini-head highlighted in purple. The cartridge is highlighted in orange (orange arrow). The microcrystal slurry droplets are ejected onto the broad face and near the downstream edge of the Kapton^®^ tape, and are aligned to the X-ray interaction region (red cross) wherein the X-ray photon path is nearly parallel to the tape face.

#### Droplet Ejection system

Previous DoT tape drive systems used acoustic droplet ejection (ADE) to dispense 2–5 nL droplets containing microcrystals onto a moving Kapton^®^ tape (Hadimioglu et al., 2016; Soares et al., 2011). To achieve lower sample consumption, we employed the piezoelectric injector (PEI) developed by PolyPico Technologies Ltd. (Cork, Ireland) (Miniature Dispensing Head) (Butryn et al., 2021). This device can dispense individual microcrystal-containing droplets in the range of 50 to a few hundreds of picolitres at a rate of up to 50 kHz. The reduction in droplet volume significantly decreases sample usage and will improve the signal-to-noise ratio but makes dehydration effects more prominent. To minimise dehydration, we positioned the ejection as close as possible to the interaction point (∼ 60 mm), with future designs incorporating a controlled humidity environment to mitigate such issues.

The droplet ejection module comprised (i) the PEI head and sample cartridge (used to hold the crystal slurry); (ii) the sample camera and the strobe LED (used to visualise ejection) iii) the wide-angle camera (used to monitor sample consumption) (Fig. 1c). For longer duration experiments, the PEI cartridge was continuously topped up with fresh samples via a fused silica capillary (with inner diameter of 150 µm) using a remote-controlled precision 250 µL syringe pump (KDS Legato× 130) sitting on a rocker to prevent sample settling in the syringe. Only the minimum amount of sample (10-20 µL) is maintained in the cartridge preventing crystal settling and clogging of the cartridge aperture.

#### Tape drive and wash-dry mechanism

The prototype belt drive utilised a brushless DC motor (MAXON) to power the conveyor belt made of Kapton^®^, 6 mm wide and 50 µm thick (kb RollerTech). The Kapton^®^ tape was treated with *Rain Repellent* (RainX) to make the surface hydrophobic thus improving drop adhesion and shape. The moving tape was cleaned after the interaction region by passing through a 3D printed module designed to pressure wash and air dry the tape before it was re-injected with sample (Fig. 1b). Although the tape drive can be operated at higher speeds, for this study we restricted the speed to 25 mm/s to ensure effective tape cleaning and sufficient residence time for microcrystals in the X-ray beam. This tape speed allows for a 720 μs maximum X-ray exposure, assuming an 8 µm crystal traversing the 10 µm X-ray beam, suitable for this application. We anticipate using a wide range of tape speeds in the final design (Table 1).

**Table 1.** Crystal exposure time to X-rays vs Tape speed.

| Linear tape speed in mm/s | Maximum time required for an 8 $\mu\text{m}$ crystal to traverse a 10 $\mu\text{m}$ X-ray beam (ms) | Dose* in kGy |
| --- | --- | --- |
| 10 | 1.8 | 31.3 |
| 25 | 0.72 | 12.5 |
| 50 | 0.36 | 6.3 |
| 100 | 0.18 | 3.1 |
| 200 | 0.09 | 1.6 |
| 500 | 0.036 | 0.6 |
| 1000 | 0.018 | 0.3 |

\*Dose calculated with RADDose-3D (Bury et al., 2018; Paithankar & Garman, 2010) based on a $8 \times 3 \times 3 \mu\text{m}^3$ CTX-M-15 crystal, and 100% transmission at 16 keV at VMXi with a flux of $2 \times 10^{13}$ ph/s at VMXi .

#### Synchronisation and X-ray data collection

Experiments at XFEL beamlines typically rely on the pulse generated by the source as their master clock for synchronisation. For these synchrotron experiments a local waveform generator (33600A, Keysight) created a “master” TTL pulse at the desired frequency up to 500 Hz (Fig. 2a) to which all other components were synchronized. This master signal was used to trigger the PEI sample ejection onto the tape while a delayed signal was used to strobe an LED (4 V white, 5 mm) to monitor the contact of the droplet with the tape. The droplets travelling on the tape to the X-ray interaction region were visualised using an on-axis viewing camera and an orthogonal camera. Only the orthogonal camera was illuminated by a strobed LED positioned downstream behind the Kapton^®^ tape and offset to prevent shadowing of the diffraction images (Fig. 1c). The strobed LED illuminated the interaction region and enabled us to synchronize the droplet arrival at the interaction region with the TTL signal by adjusting the LED delay and strobe pulse width. This adjusted TTL signal was used to trigger detector data collection. This ensured that detector readout was started just before the drop met the beam and ended just after it had passed through the interaction zone. Minimising the exposure time without a droplet in the beam will allow us to maximise the signal-to-noise and adjust the trigger point to correct for any slight alterations in ejection and tape movement. Future iterations of the device will use this technique to control a second ejection point where compounds will be mixed with the droplets on the moving tape, thus initiating chemical mixing, but was not tested during the experiment described here.

**Figure 2.**
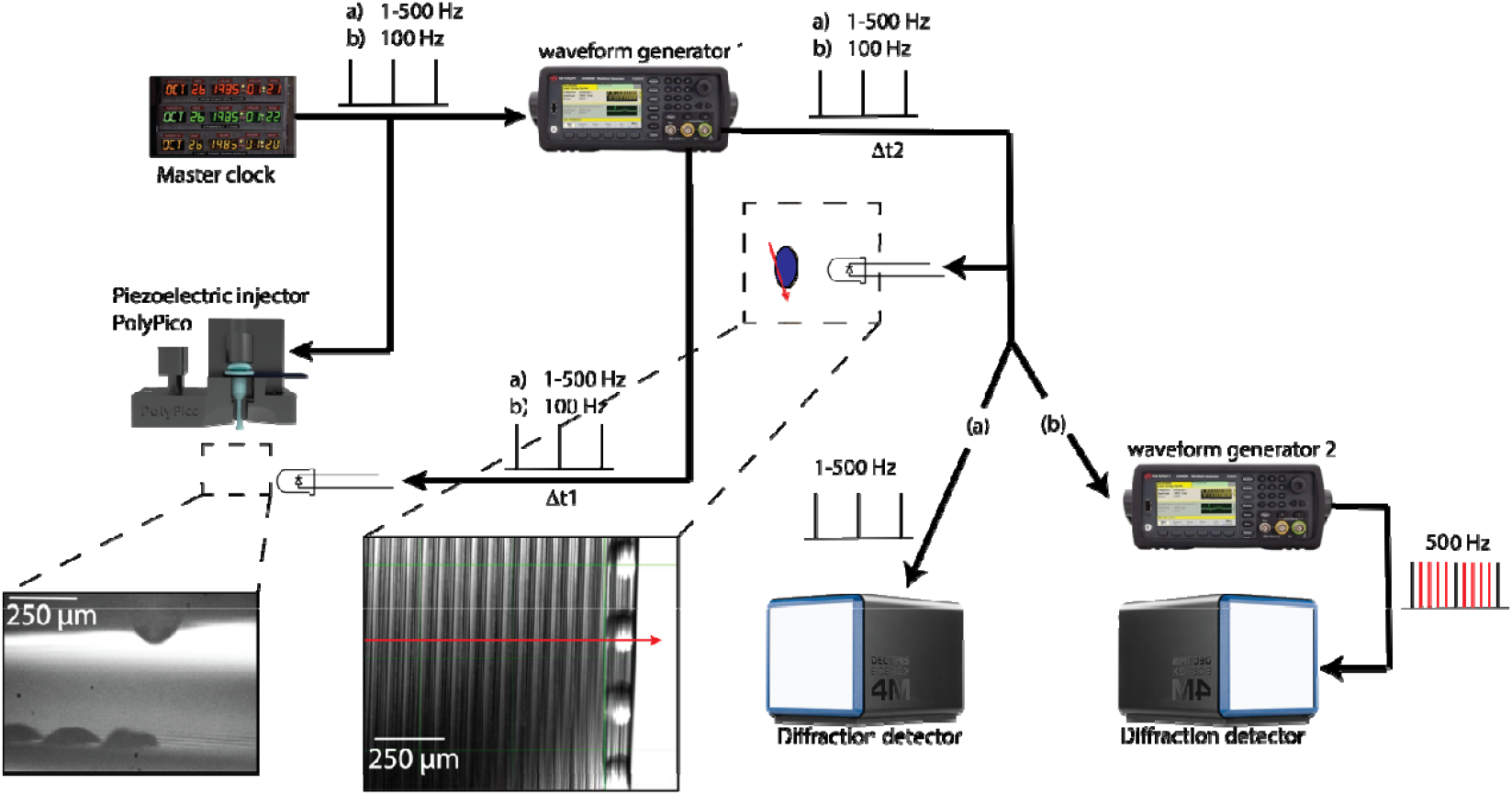
Triggering scheme of the tape drive and detector synchronisation. (a) Because synchrotron X-ray pulses are essentially continuous on the time scale of the droplet motion, a master clock is defined by a signal generator which delivers TTL pulses at the desired frequency (*e*.*g*. up to 500 Hz). This master TTL signal triggers the sample droplet ejection by the PEI. The master clock also simultaneously triggers “waveform generator 1” which will produce two TTL signals at the same frequency with delay Δt1 and Δt2, respectively. The first TTL signal triggers the strobed LED attached to the PEI; adjusting Δt1 enables the visualization and calibration of the droplet ejection onto the moving Kapton^®^ tape. The second TTL signal triggers both the strobe LED behind the tape that illuminates the X-ray – droplet interaction region, and the diffraction detector. By adjusting Δt2, the droplet position on the tape is aligned to the X-ray beam (red arrow), thereby also triggering the detector to acquiring diffraction data when droplets overlap the X-ray path. (b) Same as (a) except that the master clock runs at 100 Hz, defining the droplet ejection at 100 Hz. Another “waveform generator 2” is placed before the detector generating a 500 Hz signal. This allows ejection of droplets at 100 Hz (black lines) while acquiring images at 500 Hz (red lines).

### 2.1.3. Data collection

For data collection the linear tape speed (generated by the brushless rotating motor) used was 25 mm/s, which means that a crystal with the maximum dimension around 8 µm spent a maximum of 720 µs in a 10 µm X-ray beam, equivalent to a dose of 12.5 kGy per crystal. The PEI dispensed droplets containing CTX-M-15 microcrystals at a frequency of 100 Hz enabling the acquisition of a full dataset of >20,000 indexed images in about 30 min. The ejected droplet volume depends on several parameters such as the PEI cartridge aperture and the amplitude and width of the signal controlling the ejection. The flying droplet velocity and volume were changed by increasing both amplitude and width of the signal. In normal operation the droplet velocity before touching the tape was about 1-2 m/s. We used a PEI aperture of 100 µm to avoid clogging the cartridge with the microcrystal slurry. The amplitude (20 to 60%) and width (50%) of the ejection signal were adjusted to deliver a droplet with a volume less than 500 pL. The droplet volume was estimated by using the side view camera (see Figure 2). To ensure the success of the experiment with a sub-optimal setup (no relative humidity control) we did not try to collect data with smaller droplets (<150 µm). While the droplets were dispensed at 100 Hz the detector collection rate was 500 Hz, which corresponds to a maximum of 2 ms exposure time (Fig. 2b), of which crystals were in the beam for < 720 µs. The Eiger2X 4M detector was positioned at 230 mm from the sample position, allowing a resolution of 2.4 Å in the inscribed circle and 1.72 Å in the corner of the detector. The 16 keV X-ray beam was focused to 10 × 10 µm^2^ with a measured flux of 2 × 10^13^ photon/s.

#### 2.1.4. SSX data processing

CTX-M-15 serial synchrotron crystallography data were processed using xia2.ssx (Beilsten-Edmands et al., 2024), (Table 2). Data underwent spotfinding, indexing and integration using xia2.ssx while scaling and merging was carried out using xia2.ssx_reduce. The additional scaling parameter weighting.error_model.basic.min_Ih=300 was required to improve error modelling due to varied intensity distributions. A minor additional unit cell population was removed using unit cell clustering via the parameter clustering.threshold=5. Analysis of the indexed data versus frames collected suggested an approximate indexed hit rate of 10%. The structure was solved using PDB 7BH6 as the initial model. Structures were completed through rounds of iterative refinement in phenix.refine (Adams et al., 2010) and manual model building in Coot (Emsley & Cowtan, 2004), with validation by Molprobity (Chen et al., 2010) and Phenix.

**Table 2.**
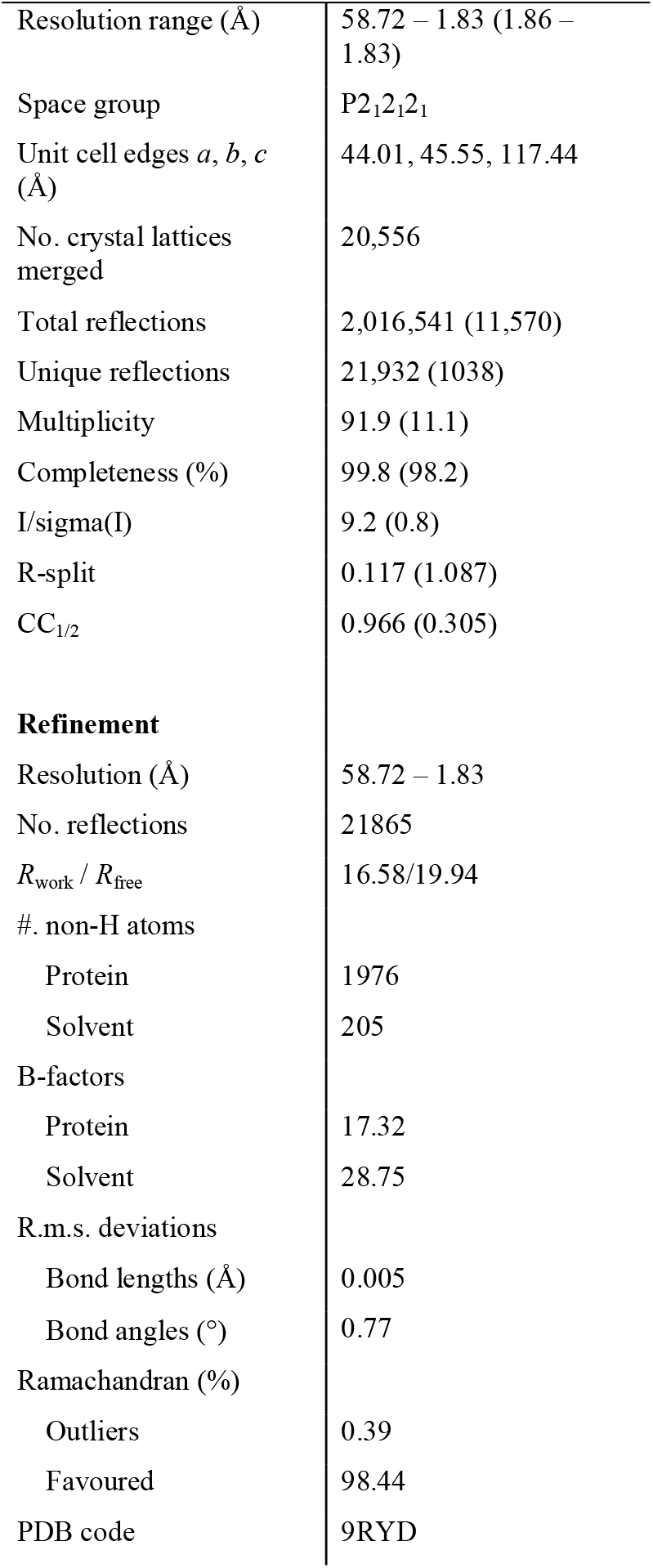
Diamond Light Source VMXi room temperature crystallographic data collection and refinement statistics for CTX-M-15 crystals.

|  |  |
| --- | --- |
| Resolution range (Å) | 58.72 – 1.83 (1.86 – 1.83) |
| Space group | P2<sub>1</sub>2<sub>1</sub>2<sub>1</sub> |
| Unit cell edges *a*, *b*, *c* (Å) | 44.01, 45.55, 117.44 |
| No. crystal lattices merged | 20,556 |
| Total reflections | 2,016,541 (11,570) |
| Unique reflections | 21,932 (1038) |
| Multiplicity | 91.9 (11.1) |
| Completeness (%) | 99.8 (98.2) |
| I/$\sigma$(I) | 9.2 (0.8) |
| R-split | 0.117 (1.087) |
| CC<sub>1/2</sub> | 0.966 (0.305) |
| **Refinement** |  |
| Resolution (Å) | 58.72 – 1.83 |
| No. reflections | 21865 |
| *R*<sub>work</sub> / *R*<sub>free</sub> | 16.58/19.94 |
| #. non-H atoms |  |
| Protein | 1976 |
| Solvent | 205 |
| B-factors |  |
| Protein | 17.32 |
| Solvent | 28.75 |
| R.m.s. deviations |  |
| Bond lengths (Å) | 0.005 |
| Bond angles (°) | 0.77 |
| Ramachandran (%) |  |
| Outliers | 0.39 |
| Favoured | 98.44 |
| PDB code | 9RYD |

### 2.2. X-ray emission spectroscopy instrumentation at VMXi

A von Hamos spectrometer to measure XES was developed in house at Diamond following similar principles to previously published designs (Fuller et al., 2017) and installed on beamline VMXi. This consisted of three components a crystal analyser array, custom built helium cone, and a 2D X-ray sensitive detector (Fig. 3). The geometry was chosen to capture the emitted fluorescence orthogonal to the main beam path and diffracted down on to a detector below the sample position. The crystal bending radius was 400 mm balancing required spectral resolution with capturing a large solid angle within the available space and minimising clashes with existing equipment. Unlike scanning spectrometers, the von Hamos geometry only focuses the X-rays horizontally, therefore the Bragg angle and associated energy vary across the vertical face of the analyser crystal. Provided the analyser crystals has sufficient height the entire spectrum can be collected simultaneously (Fig. 3). producing a vertical signal on the detector,

**Figure 3.**
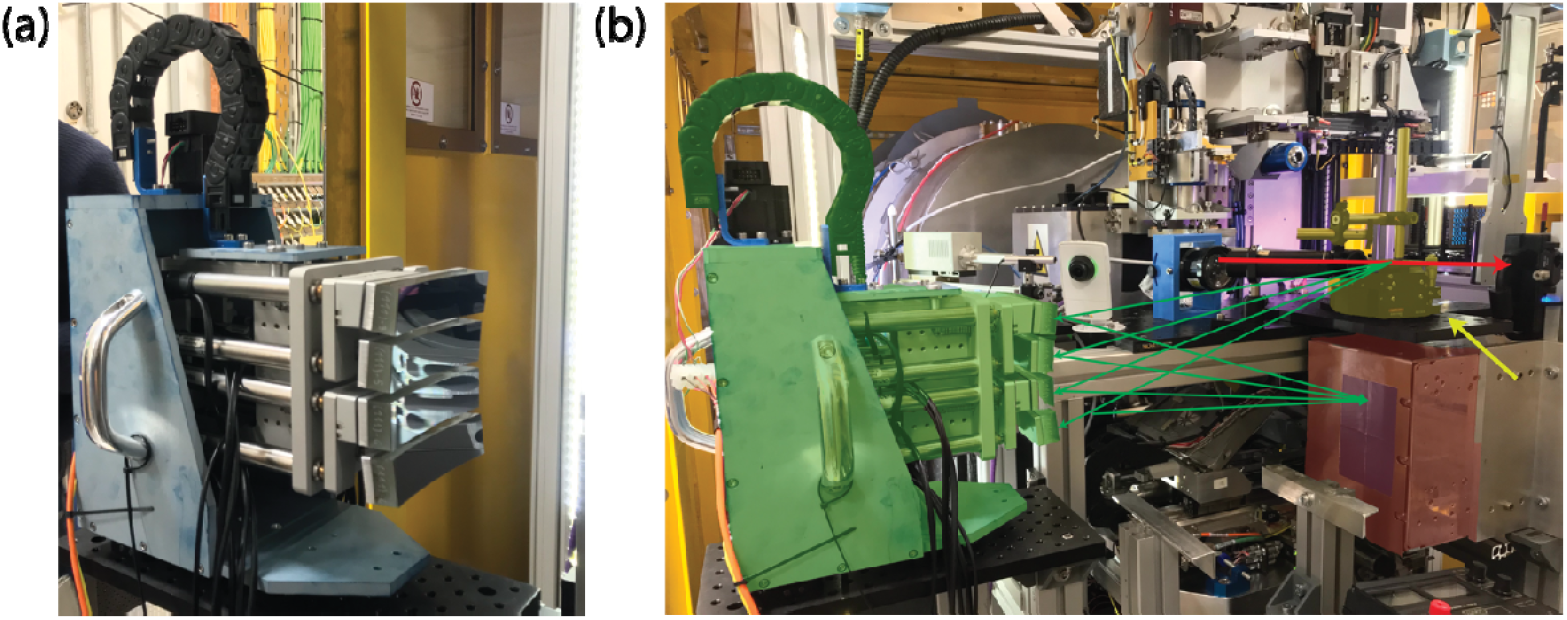
von Hamos XES spectrometer prototype installation at VMXi for Cu Kα_1,2_ measurements. A 4×1 bank of Si(111) analyser crystals with a 400 mm focal radius used to measure Cu Kα_1,2_ XES data with the Si(444) reflection. (b) Setup without helium cone and lead shielding. The XES detector (Tristan1M) is highlighted in red, the analyser crystals bank is highlighted in green, and the sample delivery system is highlighted in yellow (yellow arrow). The X-ray beam is indicated by the red arrow, and the emitted photon path is shown with green arrows.

For the proof-of-concept experiments described here we used analyser crystals arranged in a vertical block. Both Cu K□_1,2_ and Fe K□_1,2_ emission lines were collected using four Si(111) (reflection 444) or two Ge(110) crystals (reflection 440) respectively, both types with a 250 mm bending radius. Iron measurements used analyser crystals kindly borrowed from XRSTech. Each analyser crystals could be independently adjusted in focal length, pitch and yaw which was used to overlap the separate signals to maximise signal-to-noise. This adjustment was carried out by “eye” using reference metal foils or chemical solutions. For future designs we aim to incorporate automated calibration routines to ensure a consistent setup.

A simple temporary helium cone was constructed from transparent acrylic with epoxy joints. X-ray input and output windows were lined with Kapton to minimise X-ray absorption. The cone was mounted between the analyser crystals and detector and maintained at a slight positive pressure. The cone reduced the air absorption path to only 10 mm improving signal to noise (Fig. S3). Significant further improvements to the helium cone for ease of mounting and minimal helium usage are currently under investigation. The XES detector was extensively masked with lead tape (5 mm thick) to avoid contamination from direct scattering X-ray from the equipment at the beamline near the interaction region.

The X-ray detectors used were a Merlin 0.25M (Plackett et al., 2013) and a Tristan 1M (Omar et al., 2022) Timepix3 (Poikela et al., 2014) detectors for the Fe and Cu signals, respectively. Both detectors had a 55 µm vertical pixel pitch causing minimal degradation to spectral resolution. The Tristan detector is intended to be implemented for future applications and has a 113.8 x 28.3 mm active area, which has sufficient vertical height to measure multiple different emission lines simultaneously. The Merlin was used for the initial measurements as it is more compact and already integrated detector drivers with the data-collection software. As both Merlin and Tristan are based on underlying Medipix technology we anticipate no fundamental difference with photon detection. The same downstream interaction point was used as for the SSX measurements, with the extended on-axis-viewing system and re-focused beam but required a different supporting structure to mount the von Hamos crystal array and Tristan detector.

### 2.2.1. Sample preparation for XES

Iron and copper containing salts [Cu(I)Cl (229628, Merck), Cu(II)Cl_2_ (222011, Merk), K_3_Fe(CN)_6_ (10266370, Fisher Scientific) and K_4_Fe(CN)_6_ (11452378, Fisher Scientific)] were used as solids mounted between O_2_ tight foils using standard chip-less chips (Doak et al., 2018). Iron foil used for standard was purchased from Advance Research Materials Limited (FE163808).

AcNiR was purified and microcrystals prepared as previously described (Horrell et al., 2016), their size and density were assessed by visible light microscopy. Crystals of AcNiR were tested for diffraction quality at the I24 beamline (Diamond) before transfer into a high viscosity medium for extrusion. Two parts of AcNiR microcrystals slurry were mixed with three parts of monoolein (HR2-435, Hampton Research) to form the lipidic cubic phase (LCP) prior to loading in the high viscosity extruder.

### 2.2.2. Data collection

For calibration standards simple mountings were used to position the different solids samples at the focal point. Metal foils held using kinematic posts were oriented at 45° to the X-ray beam to ensure a small source point to improve spectral resolution, while also providing an escape path for the emitted photons onto the 2D Tristan X-ray detector. The Kα_1,2_ XES of the iron foil was collected with 60 s of data acquisition, and the iron and copper salts were collected with 6 s of data acquisition. Protein microcrystals were ejected into the interaction point using a high viscosity extruder and “catcher” (Doak et al., 2018). This delivery system was used due to its simplicity to install and align, albeit with higher background and not permitting synchronisation of data collection.

The LCP was extruded at a rate of 2 to 5 nanolitres per minute through a nozzle size of 75 µm inner diameter. The flow rate of the ejector was varied to ensure a stable flow, resulting in an absorbed dose range of 12-35 kGy depending on the flow rate. X-ray diffraction data (not shown) were measured simultaneously with the XES data. In-order to generate individual frames for the XES data the Tristan detector was sent the associated TTL pulse for the Xray detector readout. This could be used to turn the continuous readout data into a stack of associated images. Kα_1,2_ XES of AcNiR was collected with an acquisition time of 2400 s.

### 2.2.3. Data processing

XES data recorded either on the Merlin 0.25M or on the Tristan 1M detector were processed using in-house developed software. Data reduction and processing followed procedures as outlined in previously (Fransson et al., 2018). Data collected from the Merlin detector could be summed and reduced directly. Tristan data were processed into associated readout frames utilising the associated TTL pulse from the Eiger detector using software developed at Diamond. Tristan XES data were converted into a set of images (Hatcher et al., 2022) that matched the X-ray diffraction frame number or 30 images for foil/salt data. Hot pixels were removed with a given intensity threshold. A region of interest (ROI) around the spectrum was selected and pixel intensities along the y-axis (vertical) were summed to produce the raw spectrum. Pixel intensities in two thin bands beyond and below the ROI were also vertically summed as background and subtracted from the raw spectrum. The intensity of the background-subtracted spectra was normalised to the maximum intensity of the Kα_1_ peak. In order to calibrate the XES data Fe Kα_1,2_ spectra were measured from a Fe foil (Fig. S4), and the energy calibrated for each vertical pixel by interpolating to a known Kα_1,2_ emission lines (Thompson et al., 2009). For the Cu data, CuCl_2_ data was measured using the experimental set up reported and this data was energy calibrated using CuCl_2_ data collected on I20-Scanning at Diamond Light Source by Dr Sherwin (University of Oxford).

## 3. Results

### 3.1. Tape drive operation and delivery of droplets

We established that a moving, closed loop belt of 6 mm wide Kapton^®^, similar to that used previously in the DoT LCLS system, was the best performing option. However, because we use much smaller droplets, we needed a more precise mechanical performance, very tight motion control and highly precise ejection. Achieving the required performance necessitated the development and testing of several iterations of the tape drive system and its associated wash-dry mechanism. The version used in the present study represents an intermediate design—less precise than the final design, yet sufficiently robust to enable proof-of-concept experiments at VMXi.

First, the capability to consistently position droplets on the moving tape and synchronise this with X-ray exposures was tested. X-ray diffraction data were collected to 1.83 Å resolution from microcrystals of the CTX-M-15 extended-spectrum β-lactamase, within droplets on the tape using a linear tape speed of 25 mm/s. When the PEI was positioned 60 mm from the interaction region, it took 2.4 s for the microcrystal-containing droplet to reach the X-ray beam. In this configuration, the resulting diffraction patterns were of good quality and were successfully processed (Fig. 4a). However, when the PEI was placed more than 100 mm away, no diffraction patterns were observed, likely due to dehydration effects under a relative humidity of 20% maintained in the VMXi hutch. Consequently, the final design will include an environmental chamber that can adjust humidity levels and when necessary, also eliminates O_2_ to better than 100 PPM.

**Figure 4.**
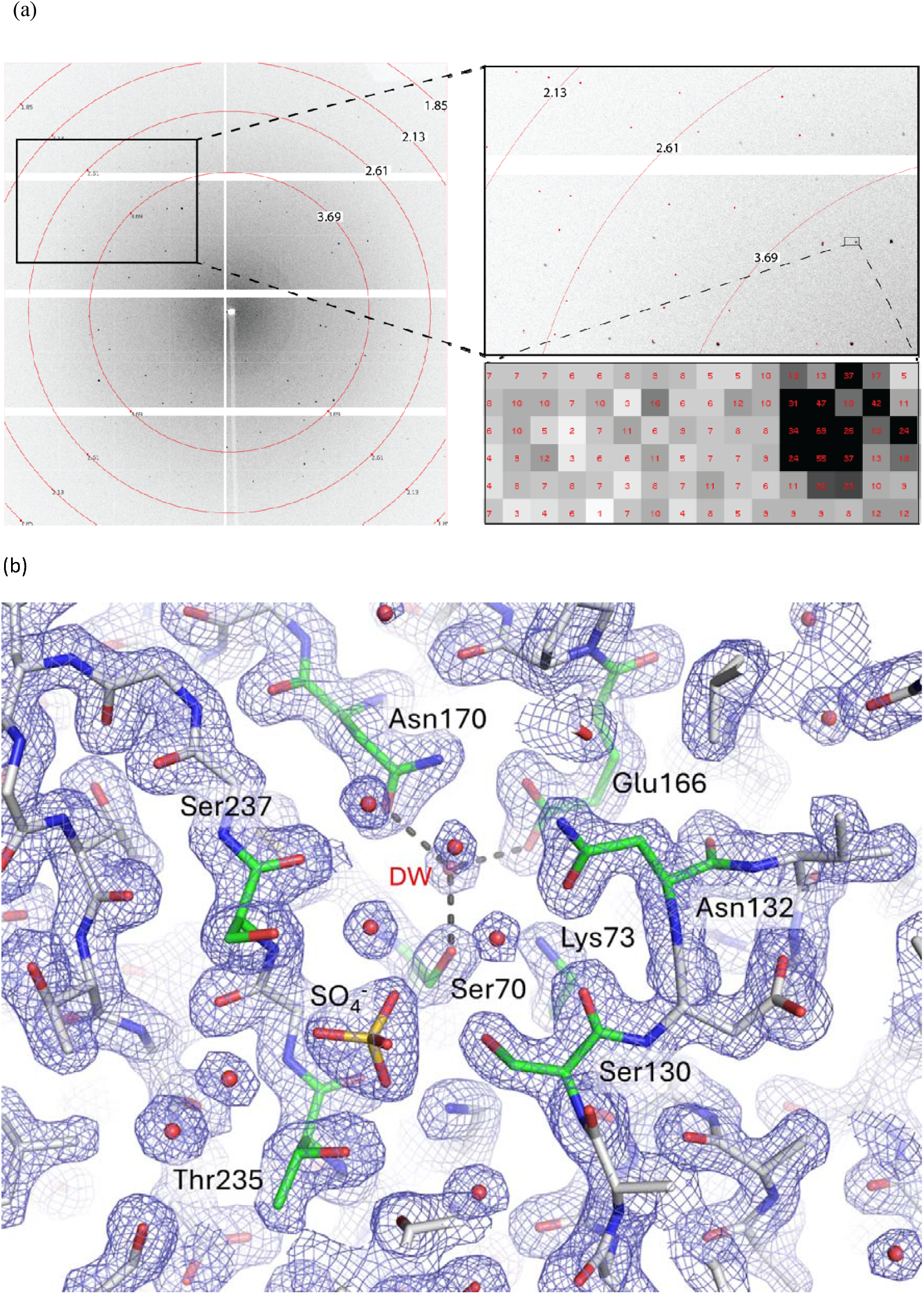
Serial synchrotron crystallography structure of serine β-lactamase CTX-M-15, originally found in *Klebsiella pneumoniae* IS53, determined using our tape drive system at VMXi: (a) Diffraction pattern measured from ∼ 8 μm microcrystals within ∼ 500 pL droplets on a tape moving at 0.25 m/s, with 2 ms total exposure time, during which crystals were in the 16 keV X-ray beam (10 ×10 µm, pink beam) for a maximum of ∼ 720 μs. Top inset: expanded regions of the detector image with an example spot profile for a ∼ 2.1 Å reflection and an indication of the background levels showing the pixel intensity across the region where the tape scattering may be observed (Bottom inset). (b) The 2Fo-Fc electron density map at 1.83 Å resolution is shown as a blue mesh contoured at 1.2σ. The carbon reading atoms of key active site residues are coloured green and labelled. Sulfate (labelled) from the crystallisation condition is bound at the active site; the catalytic, deacylating water is labelled DW and hydrogen-bonded to Ser70, Glu166, and Asn170. The figure was created in PyMol (http://www.pymol.org/).

To ensure that the X-ray beam consistently intersected with the droplets on the tape, we implemented a data collection strategy in which droplets were ejected at 100 Hz, while the detector acquired frames at 500 Hz with a 2 ms exposure time (Fig. 2b). Immediate feedback was provided by monitoring the acquired image size. When a droplet was hit by the X-ray beam the resulting image size was larger, due to the solvent ring scattering, compared to an image when no droplet was present in the X-ray path. We used this tool to move the tape as close as possible to the X-ray beam until we observed a pattern corresponding to a larger image in 1of every 5 collected (Fig. S5). By combining this approach with a Kapton^®^ tape positioned slightly off axis relative to the X-ray beam, we ensured that the beam grazed the tape surface, with even larger images being observed when the beam intersects with the tape. This geometry minimised tape-induced background scatter in the diffraction data, unlike the configuration used by the DoT which requires tape background correction tools (Fuller et al., 2017).

The main advantage of using the PEI for ejection of droplet-containing microcrystal slurry is to reduce sample consumption. Indeed, the PEI is capable of ejecting ∼150 pL droplets or less, while the ADE method (Fuller et al., 2017; Hadimioglu et al., 2016) produces droplets in the range of 2 to 5 nL depending on the transducer frequency and ejection parameters. The droplet size generated by the PEI depends on the cartridge aperture. For CTX-M-15 we used a 100 µm aperture to avoid clogging at the tip of the cartridge and to compensate the lack of relative humidity control we used higher ejection parameters than usual to create droplet volumes around 500 pL and 125 µm diameter on the tape.

Data collection took 32 min, which means that with a droplet ejection frequency of 100 Hz, we used about 100 µL of microcrystal slurry (with a density of 10^8^ crystals mL^-1^) for a high-quality dataset. The sample consumption is comparable to the fixed target approach (Horrell et al., 2021; Jaho et al., 2024). With ADE, for a similar number of indexed lattices (∼20,000) and a better indexing rate of 20% we would require at least 250 µL of the same slurry.

During the 32 min data collection, 192,000 droplets of ∼500 pL volume containing CTX-M-15 microcrystals were ejected with an average of 50 crystals in each droplet, for a crystal density of 10^8^ crystals mL^-1^. With a 10 % hit rate we indexed 20,556 lattices. In total we dispensed about 9,600,000 crystals, corresponding to 1.2 mg of protein.

The SSX data produced a clearly interpretable electron density map to 1.83 Å resolution (Fig. 4b) yielding a refined structure (Table 2) from an estimated, maximum exposure time of 720 µs (12.5 kGy) per crystal and 2 ms total X-ray time per image. With an optimised setup we anticipate reducing the sample consumption by at least a third and electronically gating the detector to better match the time each drop traverses the Xray path with commensurate reduction noise from air scatter.

### 3.2. XES data

Prototype experiments for emission spectroscopy had two principal aims, i) ensuring that anticipated Kα_1,2_ spectral changes due to different oxidation state were detectable given the experimental geometry and spectral resolution, ii) ensuring a detectable signal from protein microcrystals given the limited metal concentration within the crystal lattice, and the lower flux at synchrotrons compared to XFELs. To assess difference features the transition metals iron and copper were used as common catalytically active metals in metalloenzymes. Iron salts: potassium ferricyanide K_3_Fe(CN)_6_ and potassium hexacyanoferrate K_4_Fe(CN)_6_ were chosen to produce Fe^3+^ and Fe^2+^ signals, respectively. For copper, Copper II (Cu(II)Cl_2_) and Copper I (Cu(I)Cl) chloride were selected to produce Cu^1+^ and Cu^2+^ signals. In order to assess if sufficient signal was measurable from protein microcrystals the metalloenzyme AcNiR was used as a model system. AcNiR is easily purifiable, has a well characterised micro-crystallisation condition, contains two catalytically important copper sites, and for which catalysis can be driven by reduction in the X-ray beam, which has previously been well characterised (Horrell et al., 2016).

Iron spectra were measured from an iron foil and the ferrous salts were recorded on a 0.25M Merlin detector using two Ge(110) crystals at 250 mm working distance. Energy calibration was carried out against previously published Fe spectra (Lafuerza et al., 2020). The Fe spectra, as anticipated, produced a strong signal and a clear difference spectrum. These were in broad agreement with previous reported spectra for these compounds. Copper spectra were measured with a 1M Tristan detector replacing the Merlin, and four Si(111) crystals at 400 mm bending radius. The spectra of Cu(II)Cl_2_ and Cu(I)Cl salts, showed good signal-to-noise and a clear difference spectrum (Fig. 5b). These results demonstrate that the setup could sufficiently reproduce Fe and Cu Kα_1,2_ spectra and anticipated difference spectrum associated with the change in oxidation state.

**Figure 5.**
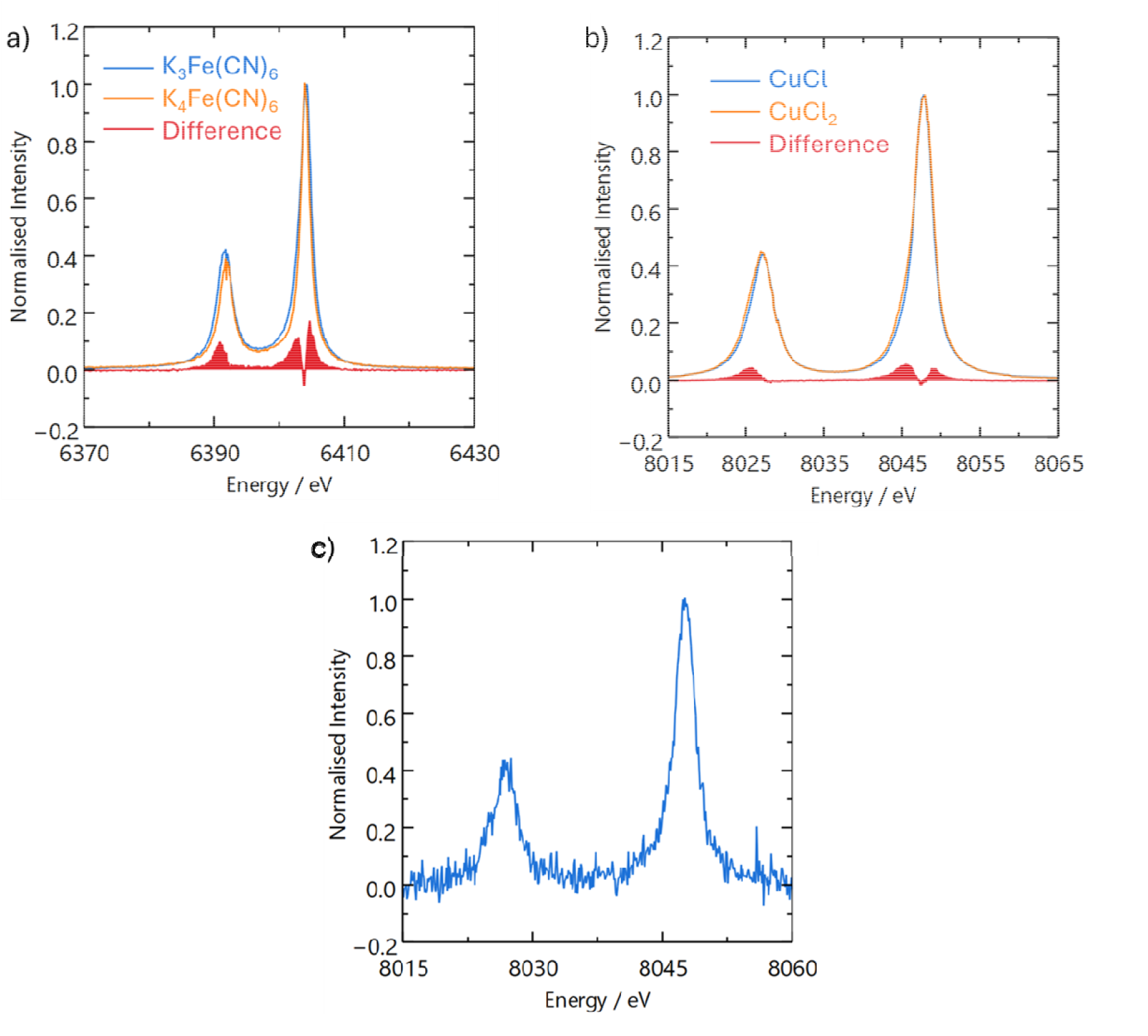
XES spectra. (a) Kα_1,2_ XES of Fe salt powder grains (K_3_Fe(III)(CN)_6_ and K_4_Fe(II)(CN)_6_) and difference spectrum; K_3_Fe(III)(CN)_6_ − K_4_Fe(II)(CN)_6_. (b) Kα_1,2_ XES of Cu salts (Cu(I)Cl and Cu(II)Cl_2_) and difference spectrum; Cu(II)Cl_2_ − Cu(I)Cl. (c) Kα_1,2_ XES of AcNiR microcrystals from the two, mononuclear Cu(II) metal centres.

Finally, we measured XES spectra from protein microcrystals using the same setup as for copper salts. Crystals of copper nitrite reductase from *Alcaligenes cycloclastes* (Horrell et al., 2018) were delivered into the X-ray beam in an LCP medium using a high viscosity extruder. The extruder is a well characterised and frequently used sample delivery method in serial crystallography, allowing us to focus on optimising the XES experiment although higher X-ray background compared to the tape drive droplets is anticipated. A clearly discernible Kα_1,2_ doublet was recorded (Fig. 5c) as an average signal collected over 2400 s that concomitantly produced a reasonable hit rate for the forward diffraction (data not shown). Comparing the calibrated spectra Centre-of-Mass (COM) to the previously collected salt spectrum did not allow assignment of the copper oxidation state in the microcrystals. Copper nitrite reductase contains two copper ions per 37 kDa monomer of the functional trimer. These copper ions have different functions and active site ligation which may lead to differences in their rate of X-ray reduction, with the resulting XES spectra likely to be an average of the signals from both copper environments and potentially also a mixed oxidation state in one or both of these.

## 4. Discussion

Our proof-of-principle experiments yielded two main outcomes (i) collection of good quality diffraction data is very possible from microcrystals dispensed onto a moving tape using ∼500 pL droplets and (ii) Cu Kα XES spectra from enzyme microcrystals delivered through the beam using a high viscosity extruder could be measured using the von Hamos setup. In both cases and despite the ongoing project in a developmental stage, good quality data were obtained. This gives us confidence that the final system will be able to deliver a strong capability, by combining these complementary measurements. Using the prototype set up described here, a crystal structure was determined using about 100 µL of microcrystal suspension. Diffraction data for CTX-M-15 were collected in about 30 minutes, producing 20,556 indexed diffraction images to 1.83 Å resolution. The resulting refined SSX structure was similar to those obtained from experiments at SACLA (drop on tape, PDB 7bh3), PAL-XFEL (fixed target, PDB 9to1), I24 (fixed target, PDB 9to5), and to a single crystal structure determined at 100 K (PDB 4HBT). The major difference between these structures is a slight movement of one of the key catalytic residues - Lys73 (particularly room temperature vs 100 K) - and of the bound water molecule responsible for breakdown of the covalent β-lactam acylenzmye complex; otherwise, the overall structures are similar with Cα RMSDs against the VMXi structure (calculated using PDBeFold) of 0.1 (SACLA, DoT, room temperature), 0.14 (PAL-XFEL, fixed target, room temperature), 0.18 (I24, fixed target, room temperature) and 0.2 (100 K structure).

For this proof-of-concept work we focused on creating a simplified tape drive mechanism to confirm that it was possible to dispense sub-nanolitre droplets of sample onto a moving tape, then present them to the X-ray beam and record diffraction data at a synchrotron beamline with sub-microsecond X-ray exposure. At VMXi the challenge of matching/synchronising the sample to the beam is slightly different to that at XFEL sources. The synchrotron beam is not pulsed (hence guaranteeing the presence of X-rays at the time the droplet crosses the interaction point). Moreover, the dose rate is much smaller and dependent on the tape velocity, which makes synchronisation a bit more difficult. For instance, at XFELs we typically observe droplet explosions and the XFEL X-ray pulse is driven by a master clock and synchronised with detectors; so, the challenge is limited to synchronising the droplet delivery to the X-ray pulse by adjusting the droplet ejection delay time relative to the master clock. Thus, the synchronisation is achieved by visual observation of droplet destruction and by observing the X-ray scattering from the droplets in the forward detector images. At synchrotrons it is not as “simple.” Unlike the XFEL pulsed X-ray beam the synchrotron X-ray beam can be considered to be constant. Therefore, to reduce unwanted background scattering caused by the constant beam, the detector and the droplet delivery into the interaction region must be synchronised. If this is not done correctly, the data are significantly limited by the weaker signal to higher noise ratio. Consequently, we used X-ray scattering from the water in the droplets that correlates with detector image file size to optimise synchronisation and maximise signal-to-noise ratio.

We expect the final design to deliver a much greater control and sensitivity, making such experiments more routine and providing a useful capability to the time-resolved structural biology community. We will use coordinated trigger signals and electronic gating of the X-ray detector to measure diffraction data only when a droplet is traversing the beam, greatly improving signal-to-noise, and allowing for shorter exposure times. It is nevertheless notable that in this study we obtained excellent quality diffraction data even without this improvement. The final system will also improve are humidity control (to avoid dehydration of samples) and implementation of controlled anaerobic conditions to permit data collection from oxygen-sensitive microcrystals that include many metalloenzymes with Fe, Cu, Mn, and/or Ni catalytic centres. As demonstrated here, reducing the droplet size from 2 – 5 nL (ADE) to between 50 – 500 pL (PEI) greatly reduces the quantity of sample required to measure a dataset, increases the signal to noise ratio, and shortens the mixing times when executing a drop-on-drop experiment. The constant but weaker X-ray source of the synchrotron compared to 10 – 50 fs pulse duration at an XFEL is also beneficial for reducing the quantity of sample needed as passing droplets through a beam that is always-on provides more opportunities for crystals to interact with the beam that may yield data.

The results with the prototype XES system are also very promising. In this proof-of-principle experiment we used a reduced solid angle (only up to 2 or 4 von Hamos analyser crystals out of the planned 16 in the final design). Furthermore, the sample delivery available at the time, LCP extruder, was far from ideal since it can be difficult to control with constant flow and produces a large X-ray background leading to much weaker signal to noise. The final version of the device will measure XES from ∼ 150 pL droplets or smaller and record the data using the newly developed Tristan 2M event counting detector. The final crystal array is planned to comprise of 4 sets of 4 analyser crystals (110 × 25 mm) with a 400 mm radius positioned to diffract back and focus the emitted fluorescence photons into an energy dispersive line on the 2D detector. The crystals are placed orthogonal to the beam path and aligned such that on the vertical plane the lattice spacing creates a discrete diffraction point on the detector depending on the energy. By selecting crystals with different lattice parameters, the device can be used to observe up to 4 separate Kα_1,2_ emission lines (planned to be used to detect Fe, Cu, Mn and Ni spectra) or, if all 16 crystals are used, we aim to be able to analyse Kβ spectra for the same elements.

Energy calibration of the Fe and Cu Kα_1,2_ XES spectra using the methods reported in 2.2.3 gave an energy axis with an energy resolution of 0.1 eV, which we anticipate being adequate for tracking changes to spin state, and oxidation state changes, across a time resolved series of XES measurements. The Kα_1,2_ XES spectra for the iron salts [K_3_Fe(CN)_6_ and K_4_Fe(CN)_6_] are congruent with literature values for both the Kα_1_ FWHM and centre of mass (COM) (Lafuerza et al., 2020) (Table 3). Successful measurements of both Fe and Cu Kα_1,2_ XES demonstrate the versatility of the von Hamos spectrometer for element-specific spectroscopic measurements by interchanging suitable analyser crystals. As expected, the Cu Kα_1,2_ XES spectrum for *Ac*NiR has a lower signal-to-noise ratio, when compared to the salt spectra; however, it is promising that the signal to noise of the protein spectrum is sufficient to calculate Kα_1_ FWHM and COM, potentially enabling tracking of changes in the oxidation state across a time-resolved reaction series. The signal-to-noise ratio for protein Kα_1,2_ XES spectra is expected to improve with the planned advances to the sample delivery system and von Hamos spectrometer. Moreover, the time-stamp capability and nanosecond temporal resolution for each pixel in the Tristan 2M detector provides opportunities to measure XES data on an orbit-by-orbit basis of the electron beam circulating in the storage ring. This enables much sharper time resolution for the spectroscopic signal compared to the current Eiger detector used in the forward direction for diffraction data. We can also easily envision a future configuration where a Tristan 10M is used for diffraction that is synchronised with the Tristian 2M for spectroscopy.

**Table 3.** XES data measured at VMXi using a von Hamos spectrometer and a Merlin 0.25M or Tristan 1M detector.

| Sample | Formal valence | Experimental $K\alpha_1$<br>FWHM /eV | COM / eV |
| --- | --- | --- | --- |
| $K_3Fe(CN)_6$ | +3 | 2.5(1) | 6399.0(1) |
| $K_4Fe(CN)_6$ | +2 | 2.0(1) | 6398.4(1) |
| CuCl | +1 | 2.8(1) | 8046.3(1) |
| $CuCl_2$ | +2 | 3.1(1) | 8044.4(1) |
| AcNiR | +2 | 2.7(1) | 8042.0(1) |

In summary, these proof-of-principle experiments demonstrate the feasibility of a synchrotron-based instrument, combining droplet on demand and XES, for routine, easy-to-use, time-resolved crystallography. To respond to the challenges encountered in this study, a device is being designed and built to collect concurrent structural and spectroscopic data in a time-resolved manner under controlled environmental conditions.

## Acknowledgements

We gratefully acknowledge the work of colleagues at Diamond beyond the author list on projects related to the development of the tape drive and XES systems. We are grateful to beamline I24 for testing of copper nitrite reductase microcrystals. We are grateful to Dr. Roberto Alonso Mori for provision of Ge(440) analyser crystals.

## Data availability

The authors declare no conflicts of interest. Coordinates and structure factors have been deposited in the Protein Data Bank with accession number 9RYD. XES spectra are made available on request.

