## Supplementary material for "Droplet on Demand Tape Drive and XES Prototypes for Time-Resolved Serial Crystallography at VMXi, Diamond Light Source": SI figures

Supporting information

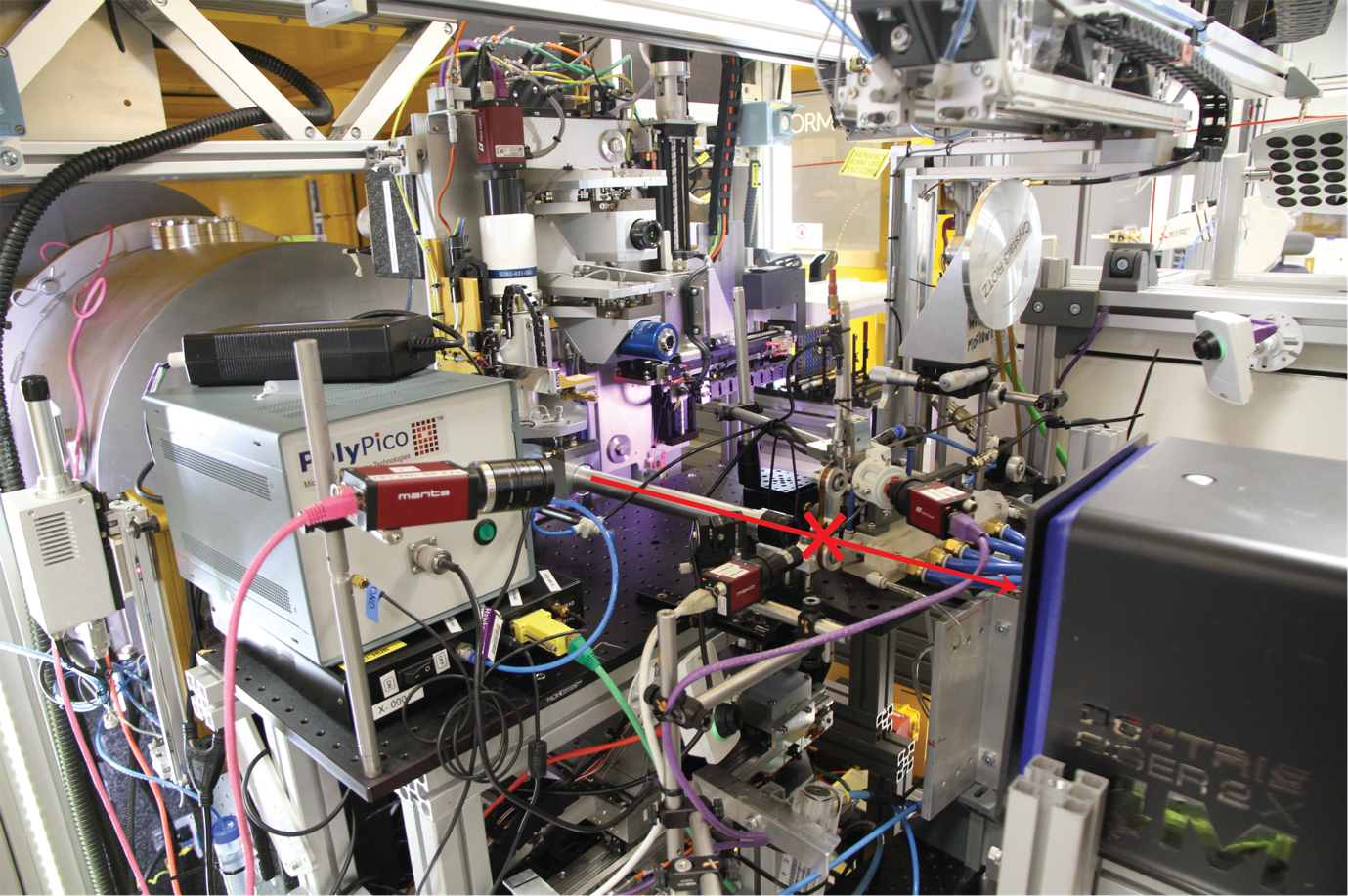

Figure S1 Tape drive setup at VMXi with the X-ray beam direction and the interaction region represented by a red arrow and a red cross, respectively.

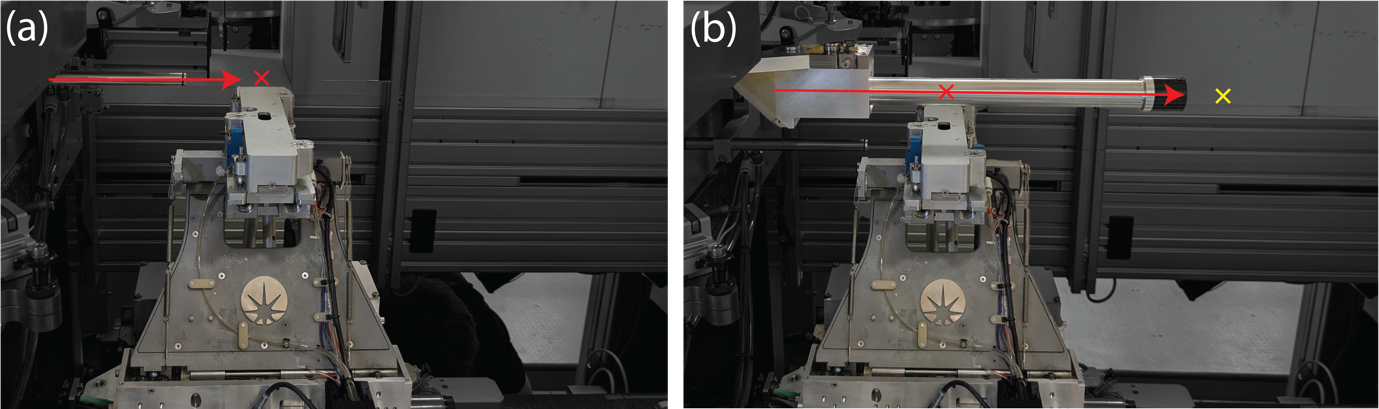

Figure S2 Interaction region with X-rays. (a) The X-rays are focussed to the position of the red crosshair when VMXi is in *in situ* mode. (b) To leave enough space to install the tape drive the beam can be refocussed 250 mm downstream (yellow crosshair). Adding an infinity-corrected objective lens allows the use of the existing VMXi beamline on-axis viewing system. The beam direction is shown by the red arrow.

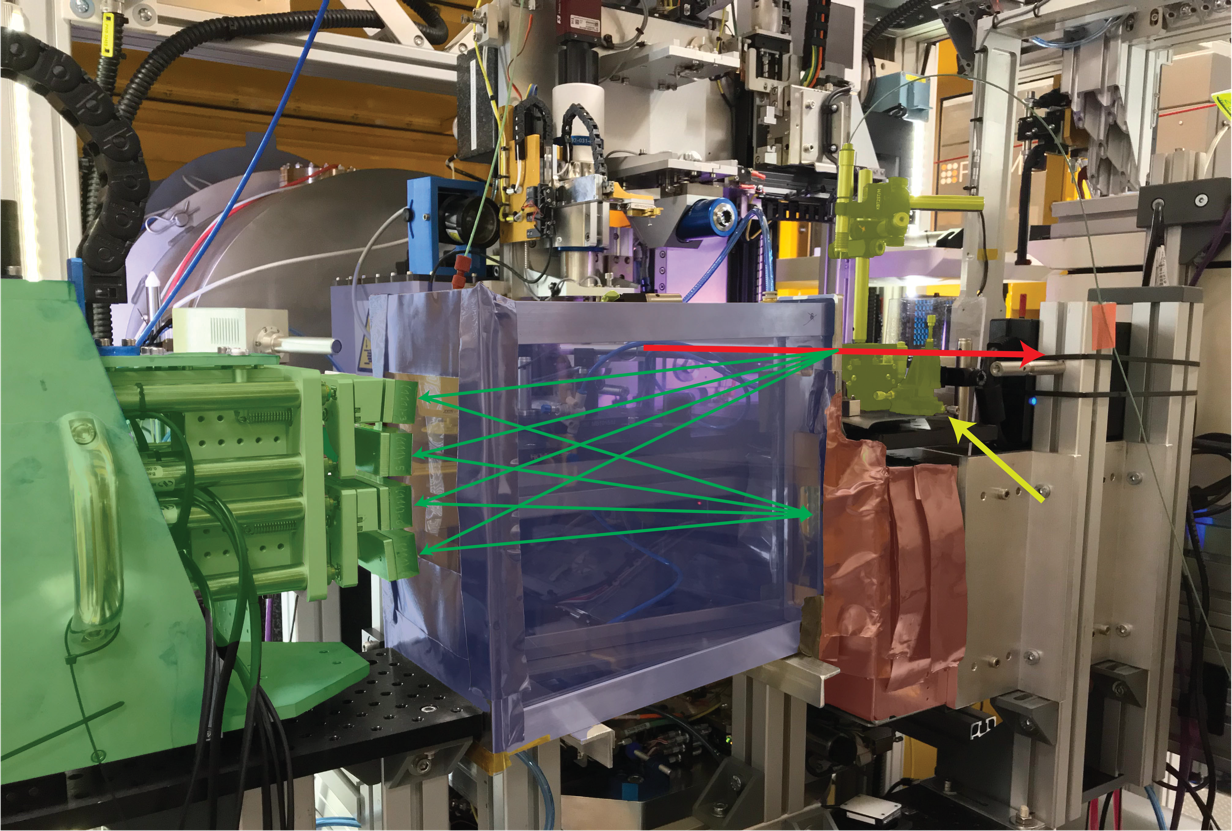

Figure S3 von Hamos XES spectrometer prototype installation at VMXi for Cu protein measurements. Setup shown with helium cone highlighted in blue and lead shielding in red shading around the XES detector (Tristan1M). The analyser crystals bank is highlighted in green, and the sample delivery system is highlighted in yellow (yellow arrow). The X-ray beam is indicated by the red arrow. The emitted photon path shown with green arrows.

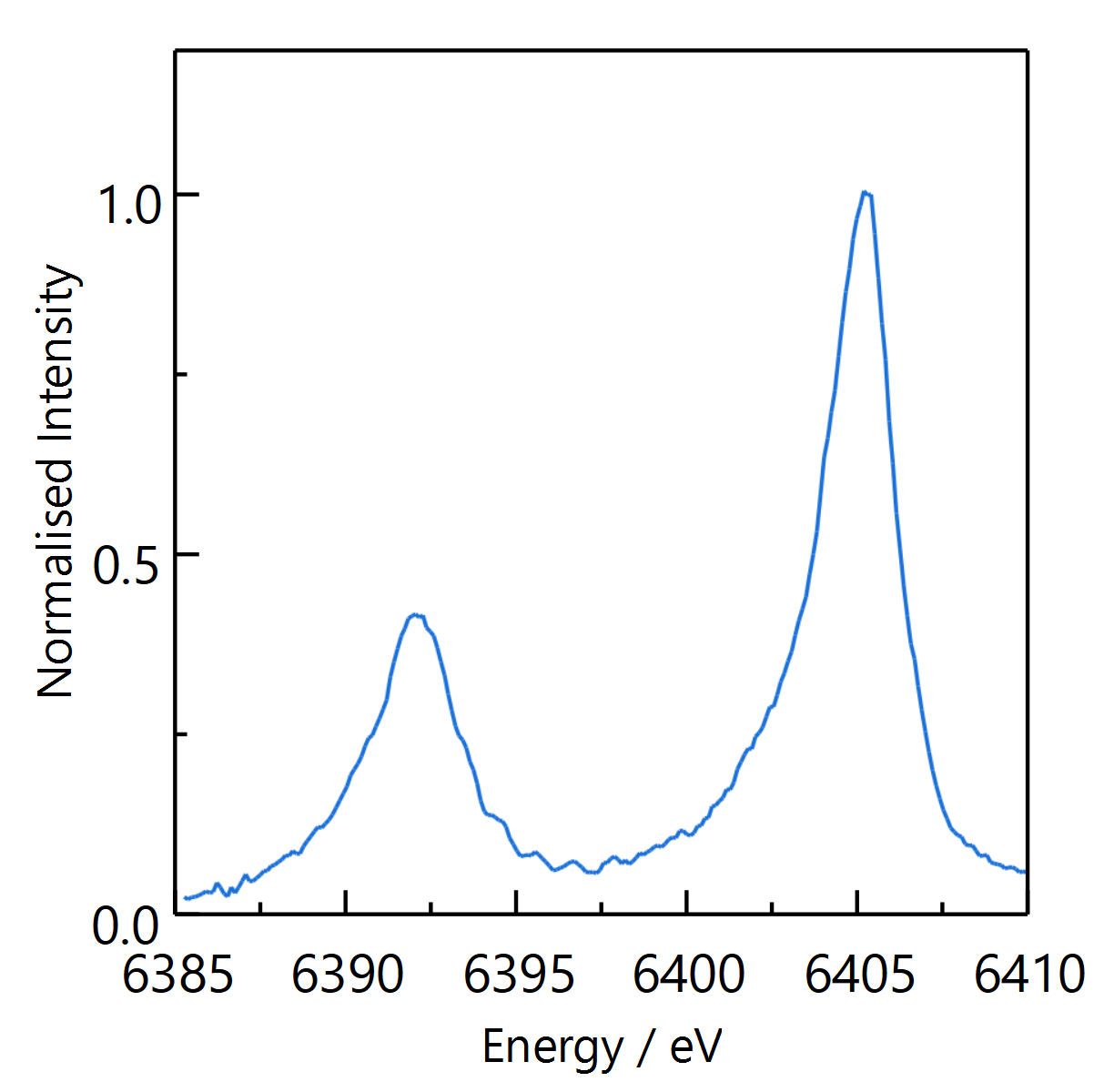

Figure S4 Kα_1,2_ XES of an iron foil measured using the von Hamos prototype set up on VMXi. A three-point smoothing filter has been applied.

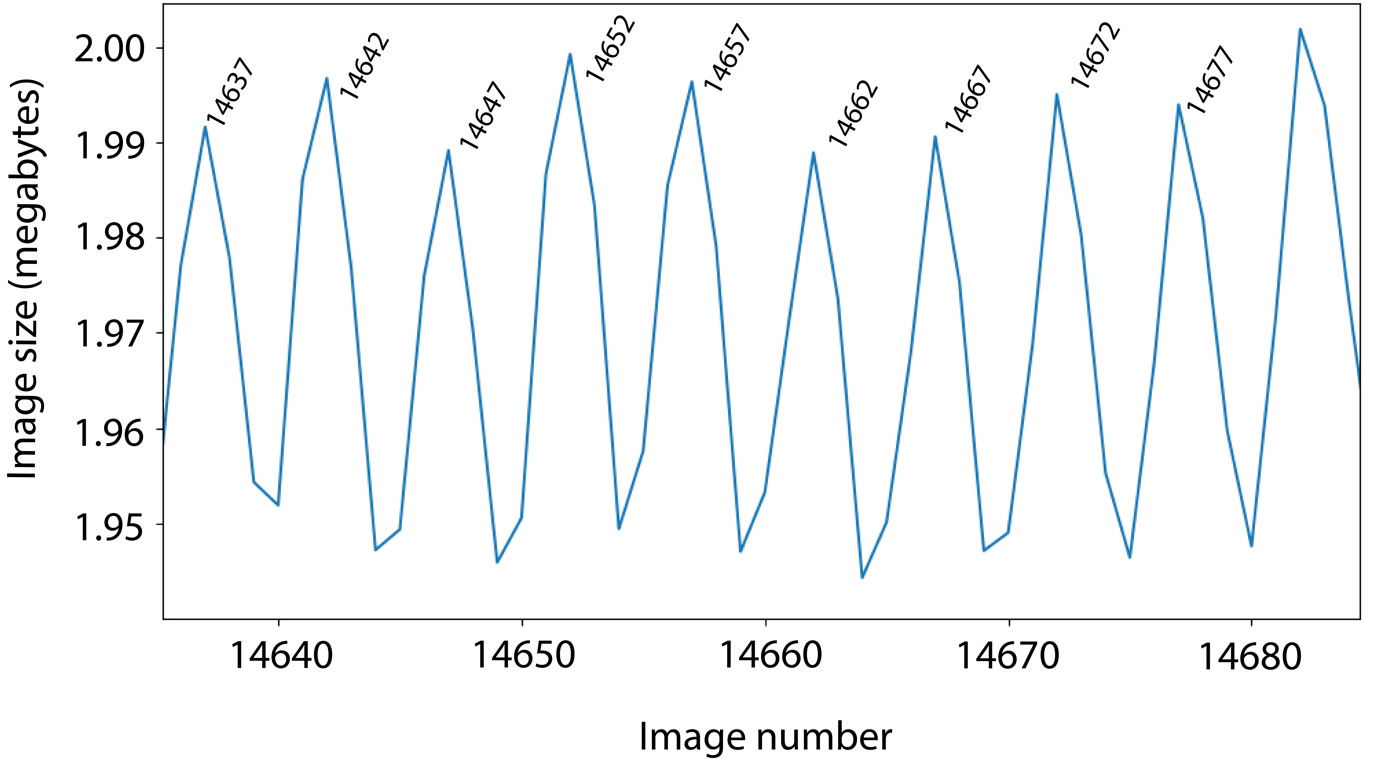

Figure S5 : Live feedback to monitor diffraction detector image size vs image number as a way to identify when a droplet is in the X-ray beam. Detector images were collected at 500 Hz but droplets were deposited onto the tape at 100 Hz. The image size is larger if/when a droplet is in the beam due to water scattering at low *q* regardless of whether or not the image also contains diffraction. A file size peak is observed every 5 images, which correlates with when a droplet is in the beam and diffraction if a crystal also intersects the beam. This serves as a useful proxy to aid synchronisation of droplets and diffraction data collection.
